# Nonlinear sensitivity to acoustic context is a stable feature of neuronal responses to complex sounds in auditory cortex of awake mice

**DOI:** 10.1101/2023.04.22.537782

**Authors:** Marios Akritas, Alex G. Armstrong, Jules M. Lebert, Arne F. Meyer, Maneesh Sahani, Jennifer F. Linden

## Abstract

The perceptual salience of a sound depends on the acoustic context in which it appears, and can vary on a timescale of milliseconds. At the level of single neurons in the auditory cortex, spectrotemporal tuning for particular sounds is shaped by a similarly fast and systematic nonlinear sensitivity to acoustic context. Does this neuronal context sensitivity “drift” over time in awake animals, or is it a stable feature of sound representation in the auditory cortex? We used chronically implanted tetrode arrays in awake mice to measure the electrophysiological responses of auditory cortical neurons to spectrotemporally complex, rapidly varying sounds across many days. For each neuron in each recording session, we applied the nonlinear-linear “context model” to estimate both a principal (spectrotemporal) receptive field and a “contextual gain field” describing the neuron’s nonlinear sensitivity to acoustic context. We then quantified the stability of these fields within and across days, using spike waveforms to match neurons recorded in multiple sessions. Contextual gain fields of auditory cortical neurons in awake mice were remarkably stable across many days of recording, and comparable in stability to principal receptive fields. Interestingly, there were small but significant effects of changes in locomotion or pupil size on the ability of the context model to fit temporal fluctuations in the neuronal response.

We conclude that both spectrotemporal tuning and nonlinear sensitivity to acoustic context are stable features of neuronal sound representation in the awake auditory cortex, which can be modulated by behavioral state.

## Introduction

Are sensory receptive fields of neurons in auditory cortex of adult animals fundamentally stable properties? Decades of research has shown that auditory cortical receptive fields in adult animals can be altered by behavioral training or by shifts in auditory attention (for reviews see ***Weinberger, 2007; Fritz et al., 2007; Irvine, 2018***). However, the long-term baseline stability of auditory cortical receptive-field structure in adult animals is less well studied — especially for nonlinear features of the receptive fields, such as sound combination sensitivity and other forms of modulation by acoustic context. Here we analyze the long-term stability of nonlinear context sensitivity as well as spectrotemporal tuning in auditory cortical receptive fields of awake mice, using chronic electrophysiological recording and nonlinear stimulus-response function estimation.

The stability of sensory cortical response properties in awake animals has recently become a hot topic in debates about the nature of “representational drift” (***Clopath et al., 2017; Driscoll et al., 2022; Marks and Goard, 2021***). In the auditory cortex, long-term two-photon calcium imaging studies in awake mice have reported “representational drift” in population responses to sound stimuli (***Kato et al., 2015; Chambers et al., 2022; Aschauer et al., 2022***). Notably, this “drift” appears to arise primarily from changes in whether individual neurons respond to their preferred stimuli, rather than changes in their stimulus preferences when responsive (see for instance Supplementary Figure 5 in ***Chambers et al., 2022***). However, the slowness of the calcium signal makes it difficult to reconstruct details of auditory cortical receptive fields, and therefore difficult to determine the extent to which spectrotemporal tuning and combination sensitivity might remain consistently stable in individual neurons when they are responsive. Sound patterns that evoke the same response from a neuron on the slow timescale of calcium signalling could evoke different responses measured at the fast timescale of neuronal spiking. Thus, previous calcium imaging studies in auditory cortex have not resolved questions about whether auditory cortical receptive fields in adult animals remain stable across days when measured with the millisecond temporal resolution most relevant to auditory perception.

Previous electrophysiological studies have reported that spectrotemporal tuning of auditory cortical neurons can remain stable for many hours — but to the best of our knowledge, no studies have investigated the long-term stability of nonlinear sensitivity to acoustic context. ***Elhilali et al. (2007***) used reverse-correlation techniques to obtain repeated estimates of linear spectrotemporal receptive fields (STRFs) from neuronal responses to complex sounds in awake, passively listening ferrets, and found that STRF structure of individual neurons remained relatively stable across many hours of recording. Similarly, ***Grana et al. (2009***) reported that STRFs recorded from neurons in field L (avian auditory cortex) were stable for hours in awake, passively listening songbirds. Other electrophysiological studies focusing on more basic measures of spectrotemporal selectivity, such as frequency tuning and spike timing statistics, have suggested that neuronal response properties might be stable for days or weeks in awake animals implanted with electrode arrays (***Williams et al., 1999; Witte et al., 1999***). How stable are spectrotemporal receptive fields of auditory cortical neurons in awake animals across days or weeks? And how stable are nonlinear features of auditory cortical receptive fields, such as sound combination sensitivity and modulation by acoustic context? To address these questions, we analyzed the stability of auditory cortical receptive-field structure in awake, passively listening mice across days of chronic electrophysiological recording, using the nonlinear-linear “context model” (***Williamson et al., 2016***) to estimate both spectrotemporal tuning and contextual sensitivity of auditory cortical neurons from their spiking responses to complex sounds. Unlike the STRF and commonly used linear-nonlinear models of auditory cortical responses (reviewed in ***Meyer et al., 2017***), the context model allows for nonlinear integration of spectrotemporal elements within a complex stimulus (for example, nonlinear forward suppression or two-tone interactions). The context model includes both an STRF-like principal receptive field (PRF), with dimensions of sound frequency and time preceding the neuronal response, and a contextual gain field (CGF), with dimensions of frequency offset and time offset for sound combinations that modulate input gain. The PRF represents the spectrotemporal tuning of the neuron, while the CGF represents its contextual sensitivity — i.e., how the responsiveness of a neuron to a tonal element within a complex sound is affected by the acoustic context in which that element appears. Hence, the CGF captures suppressive or facilitatory effects of sound combinations, which modulate the gain of the neuron’s response to each sound element falling within the PRF. Estimation of PRF and CGF parameters from neuronal responses to a complex sound is an experimentally efficient and relatively stimulus-agnostic means of determining both spectrotemporal tuning and nonlinear sensitivity to acoustic context (***Williamson et al., 2016; Meyer et al., 2017***).

We report that neuron-specific patterns of nonlinear contextual sensitivity as well as spectrotemporal tuning remain stable across multiple days in awake, passively listening mice. We also observe significant but very small effects of changes in locomotion or pupil size on the ability of the context model to fit temporal fluctuations in auditory cortical responses. These results suggest that auditory cortical neurons can maintain consistent receptive fields for many days, despite some modulation by spontaneous behavioral state. Moreover, the findings indicate that nonlinear tuning to acoustic context is a robust and remarkably stable feature of the neural code in the awake auditory cortex.

## Results

We chronically implanted male CBA/Ca mice with multi-tetrode arrays, using a tangential approach to the auditory cortex. We used the tetrodes to record extracellularly from auditory cortical neurons in awake, head-fixed mice, while also measuring running behavior and pupil diameter, while the animals listened passively to noise bursts, tone pips and dynamic random chord (DRC) stimuli (Figure 1). Two DRC stimuli, each consisting of 15 continuous repetitions of a 45-s-long DRC trial, were presented within each recording session. We conducted multiple recording sessions at each recording site, repeating exactly the same stimulation and recording protocol on different days. In 4 mice we obtained high-quality auditory cortical recordings from multiple recording sites across at least 5 days for each site, which could be used to assess stability of neuronal responses over time.

**Figure 1.**
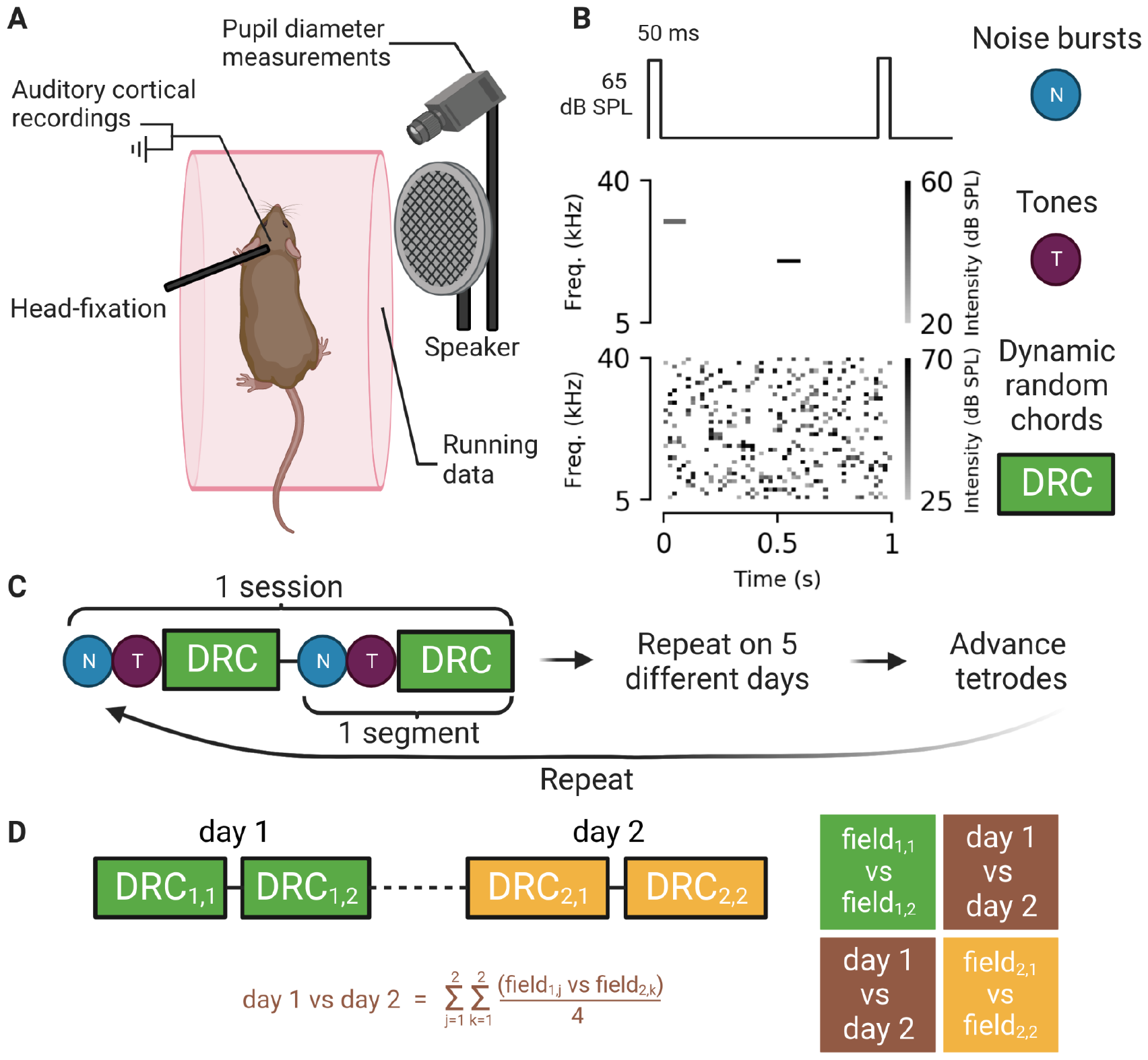
Experimental setup, auditory stimuli, recording strategy, and stability assessment. **A**. Illustration of experimental setup. Single-unit and multi-unit recordings were obtained from the auditory cortex of awake mice using a chronically implanted 8-tetrode array. Mice were head-fixed but were able to run on a rotating cylinder. Simultaneous neuronal recordings and measurements of running speed and pupil diameter were obtained during repeated presentations of noise bursts, tone pips, and dynamic random chord (DRC) stimuli. Responses to noise bursts and tone pips were used to identify core auditory cortical areas. Responses to DRC stimuli were used to estimate contextual gain fields (CGFs) and principal receptive fields (PRFs) using the context model. **B**. Schematic illustrations of noise bursts (top), tones (middle), and DRC excerpt (bottom). A full DRC stimulus lasted 675 s, and consisted of 15 continuous repetitions of a 45-s-long sequence of 20-ms random chords. **C**. Schematic representation of the experimental design. Recordings were obtained from the same site for multiple days before tetrodes were advanced to sample new sites. Note the repetition of the full sequence of stimulus presentations (1 segment) within each session. **D**. Conceptual illustration of methodology used for assessing stability of the context model fits. On each day of recording, the full DRC stimulus was played twice, once in each segment (upper left). CGF or PRF estimates from different days and/or different segments (field_*j,k*_, where *j*=day and *k*=segment) were compared both within and between sessions to obtain a similarity matrix (right). Within-session similarities are on the diagonal (in green and yellow) and the average estimates of the across-session similarities (lower left) are on the off-diagonals (in brown).

Recordings were spike-sorted to distinguish single units from likely multi-units (see Materials and Methods). Recording sites in core auditory areas were then identified as those producing unit recordings with significant and short-latency (≤ 20 ms) responses to tone pips (Supplementary Figure 1). The core auditory dataset (314 single units, 199 multi-units) included units that either directly fulfilled these criteria or were recorded on the same tetrode at the same time (and hence at the same location) as another unit that fulfilled the criteria.

We matched neuronal recordings obtained over multiple different days by analyzing spike wave-form similarity in tetrode recordings. To do so we customized a waveform-matching technique introduced by ***Tolias et al. (2007***), which quantifies the difference between two sets of spike wave-forms using two metrics, *d*_1_ and *d*_2_. The former measures differences in shape between spike wave-forms and the latter measures differences in scale. We improved upon the original approach by establishing a null distribution (in dimensions of *d*_1_ and *d*_2_) reflecting differences between definitively non-matched units, recorded using the same tetrode but from sites spaced at least 250 microns apart (Figure 2A). We used parameters of this null distribution to distinguish likely from unlikely matched pairs in the experimental dataset, which consisted of spike waveforms recorded using the same tetrode on different days at the same recording site (Figure 2B–C; see also Materials and Methods).

**Figure 2.**
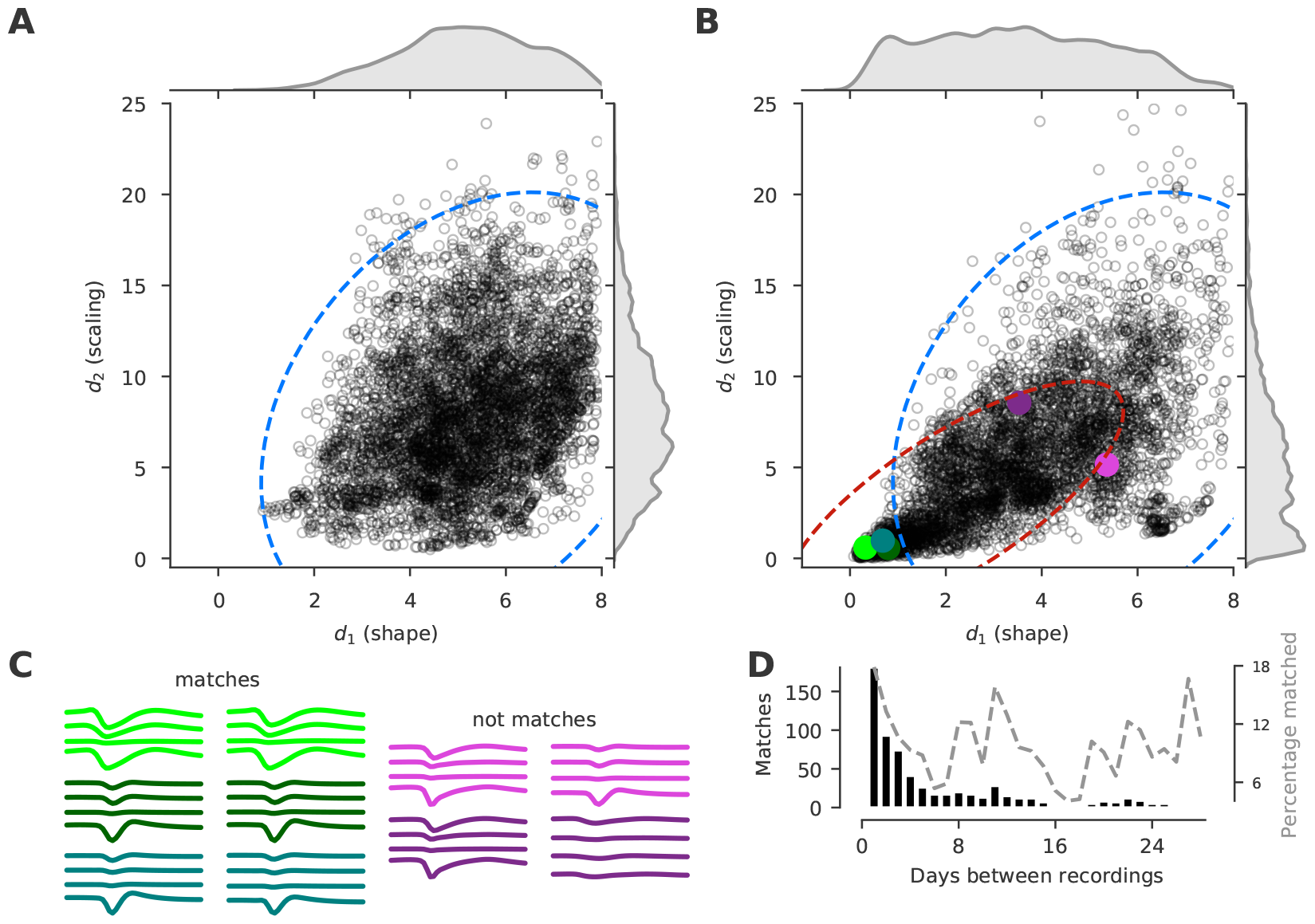
Spike waveforms matched across multiple days using pairwise waveform distances. See text for explanation of waveform distance metrics *d*_1_ and *d*_2_. **A**. Null distribution. Scatterplot shows (*d*_2_, *d*_1_) spike waveform distances for pairwise comparisons (n=6574) between spike waveforms for unit recordings known to be non-matched (obtained using the same tetrode but from sites located at least 250 microns apart). The ellipse represents the 99% confidence interval (CI) for the null distribution, estimated by fitting a 2D Gaussian to the data. Marginal distributions were obtained using kernel density estimation. **B**. Experimental distribution. Scatterplot shows (*d*_2_, *d*_1_) spike waveform distances for pairwise comparisons (n=5594) between spike waveforms for unit recordings obtained using the same tetrode on different recording days at the same recording site. A Gaussian mixture model was fitted to the experimental data using the Expectation-Maximization (EM) algorithm with two clusters. One of the clusters was fixed to the null distribution estimated in A. Ellipses show the 99% CIs for the null (blue) and the experimental (red) distributions. We conservatively defined a waveform pair to be “matched” (i.e., likely to be coming from the same unit) if the waveform distance fell within the experimental but outside the null 99% CI. Colored dots correspond to the example matched and non-matched waveform pairs shown in C. **C**. Examples of spike waveform pairs. The pairs in the first two columns were identified as matches, whereas those in the latter two columns were not. **D**. Number of matches as a function of the temporal separation between the two recordings. Dotted gray line shows percentage of total comparisons which were matches. Note that the number of waveform pairs identified as matches was highest for recordings occurring 1–4 days apart, but this was primarily because the number of pairwise waveform comparisons was highest for recordings occurring a small number of days apart. The *percentage* of waveform comparisons producing a match could be just as high for recordings made weeks apart as days apart, indicating that prolonged tetrode recordings from the same site could be stable.

We obtained 637 matches, some made between units recorded as long as 3 weeks apart. As expected, most waveform matches were made across 1–4 elapsed days (Figure 2D, histogram), because most recordings from the same site were obtained across 5 consecutive days. However, some of the intervals between recordings at the same site were much longer, and we saw no evidence for an overall decline in the fraction of waveform matches at longer intervals. For recordings separated by weeks, the percentage of waveform comparisons producing matches could be as high as for recordings separated by 1–4 days (Figure 2D, dotted line).

### Neuronal sensitivity to acoustic context in awake mice conserves main features seen in anesthetized mice

For consistency with a previous study of neuronal CGF structure in anesthetized animals (***Williamson et al., 2016***), we included in the context model analysis all units in the core auditory dataset for which the signal power in the neuronal response to the DRC stimulus was at least one standard error greater than zero (a total of 142 single units and 127 multi-units). Signal power is the stimulus-dependent power in the neural response — i.e., the portion of the temporal variability in the response that is preserved from trial to trial, and that is at least in principle predictable from a stimulus-response function model. As in previous work (***Sahani and Linden, 2002b; Ahrens et al., 2008; Williamson et al., 2016***), we defined noise power as the remaining, stimulus-independent part of the response, encompassing any variability that is not repeatable across identical trials. Context model parameters can be estimated effectively for any neuronal response with significantly non-zero signal power, regardless of noise power. Nevertheless, the signal-to-noise power ratio (SNR) provides a useful quantitative index of selectivity for the DRC stimulus in the recorded population. Figure 3A shows the SNR values for all units in the dataset, and Figure 3B–F provides examples of DRC responses for units with different SNRs. Note that both single-unit and multiunit recordings yielded DRC responses with SNRs spanning the entire SNR range observed in the population.

**Figure 3.**
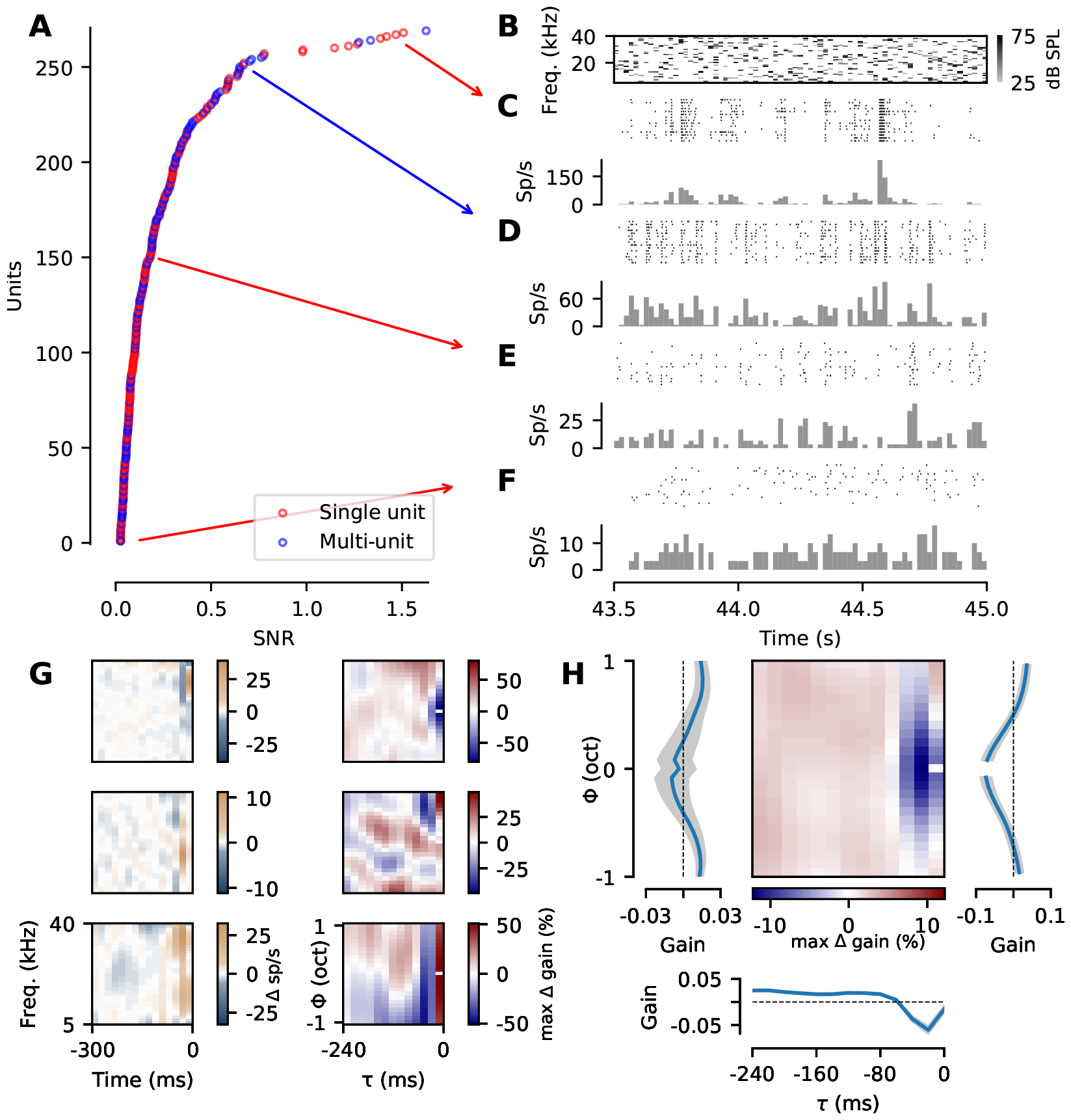
Neuronal responses to the DRC stimulus used to estimate PRF and CGF structure in awake mice. **A**. Signal power normalized by noise power (SNR) for neuronal responses to the DRC stimulus, for all units that qualified for further analysis given our selection criteria (see text). Units are sorted in order of ascending SNR. Single units are shown in red, multi-units in blue. **B**. Spectrographic reresentation of the final 1.5 s of the 45-s-long DRC stimulus used. Each shaded rectangle represents a 20-ms tone pulse, with darker shades corresponding to louder tone pips (see colorbar). **C-F**. Trial-by-trial spike rasters (top) and histograms of spiking rate (bottom), describing the responses of four example units to the DRC excerpt in B. Histogram bins are aligned with the 20-ms chords of the DRC. Units were taken from a point in the distribution in A indicated by the arrows. Time is shown relative to the beginning of the stimulus for the trial. **G**. Example PRFs (left) and CGFs (right) for three different units (each row is one unit). Yellow and cyan areas in the PRFs represent excitatory and inhibitory regions of the time-frequency receptive field respectively. In the CGFs, axes are time offset and frequency offset relative to a “target” tone represented by the notch at *τ*=0 and *ϕ*=0, which can be any tonal element in the DRC stimulus. Red and blue areas in the CGF indicate amplifying or dampening effects (respectively) of acoustic energy at that relative position on the gain of the neuron’s response to a target tone. In other words, the CGF depicts modulation of neuronal responsiveness by sound combinations, as a function of time and frequency differences between the tonal elements in the combinations. **H**. Average CGF across all units and animals (center). For units recorded across multiple days, we included in this average a single CGF estimated from all the available data for the unit. Line plots along margins show: (left) gain profile as a function of frequency offset between tone pips, averaged across time offsets; (bottom) gain profile as a function of time offset between tone pips, averaged across frequency offsets; and (right) gain profile as a function of frequency offset for the 0–20-ms time-bin alone (i.e., for near-simultaneous tone pips). Error bars indicate standard error of the estimated population means.

To examine features of contextual sensitivity in auditory cortical neurons of awake mice, we first fit a single context model for each unit, pooling all the DRC responses recorded from that unit across multiple days of recording in the awake animal. Procedures for estimating the PRF and CGF parameters in the context model are described in Materials and Methods. Figure 3G depicts example PRFs and CGFs from three units, and Figure 3H shows the average CGF across all units and animals. This average CGF illustrates the most common features of contextual sensitivity in auditory cortical neurons of awake, passively listening mice, and it is consistent with that previously reported by ***Williamson and Polley (2019***) specifically for neurons in layers 5 and 6 of awake mouse auditory cortex.

Notably, contextual sensitivities of auditory cortical neurons in awake passively listening mice were also qualitatively similar to those previously described in anesthetized mice (***Williamson et al., 2016***). Like the average CGF for anesthetized mice shown in that reference, the average CGF for awake mice (Figure 3H) exhibited (i) narrowband delayed suppression (blue region centred on zero frequency offset and extending over negative time offsets) and (ii) near-simultaneous broadband facilitation (red areas at zero time offset and large frequency offsets on either side of the target tone). However, the narrowband delayed suppression observed previously in anesthetized mice peaked at and extended to longer time offsets than was observed in awake mice here (and by ***Williamson and Polley, 2019***).

### Neuronal sensitivity to acoustic context is stable across days of recording in awake mice

We next examined whether both spectrotemporal tuning and contextual sensitivity of auditory cortical neurons were stable over time in awake mice. To do so, we compared different estimates of PRFs or CGFs obtained from the same unit within and across recording sessions, using the spike-waveform-matching technique described previously to track units across multiple days of recording at the same recording site. Within-session comparisons of repeated PRF or CGF estimates provided a measure of short-term test-retest reliability, while across-session comparisons allowed us to measure long-term stability of spectrotemporal tuning and contextual sensitivity.

As demonstrated by the examples in Figure 4A-C, the structure of both PRFs and CGFs was often remarkably consistent across recording days. We calculated the normalized dot product (field correlation) between repeated PRF or CGF estimates obtained for the same unit to assess consistency of neuron-specific structure. Correlation between repeated PRF or CGF estimates for the same unit could be nearly as high across days as within recording sessions (Figure 4D-F).

**Figure 4.**
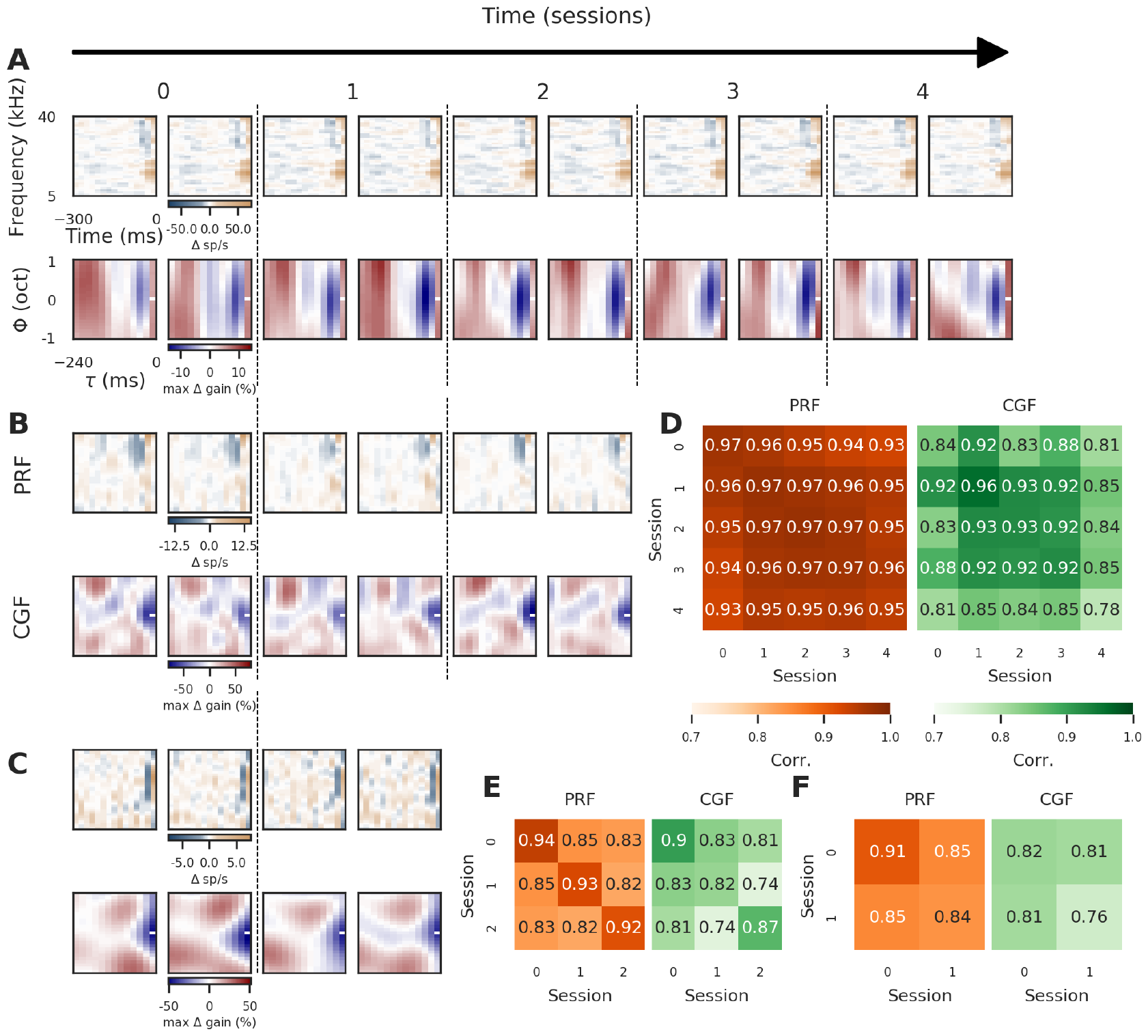
Examples of quantification of PRF and CGF stability across recording days. **A-C**. Example PRF (top row) and CGF (bottom row) pairs for neurons matched across recording sessions. The within-session repetition of the 675-s-long DRC run allowed us to estimate two PRFs and two CGFs for each session. For each example, PRFs are identically scaled to the maximum change in firing rate shown in the PRF colorbar. CGF weights at each value of (*ϕ, τ*) represent the change in gain induced in the response to a sound at the (0,0) notch point if one of the loudest tones of the DRC were to fall at the corresponding (*ϕ, τ*) location (colors correspond to gain change shown on the CGF colorbar). Like PRFs, CGFs are identically scaled within and across sessions for each example. Time runs from left to right and is in recording sessions conducted on separate but not necessarily consecutive days; numerals across the top of panel A indicate number of recording sessions following the initial session. Note the remarkable consistency of both CGF and PRF structure, which is nearly as high across days as within sessions. **D-F**. Heatmaps showing the normalized dot product (i.e., field correlation) between the PRFs (orange) or between the CGFs (green) shown in A-C, respectively. Diagonals indicate the within-session comparisons, off-diagonals the across-session (i.e., across-day) comparisons. Higher values indicate higher correlation in structure. The correlation color scale was set to 0.70–1.00 (rather than 0.00–1.00) to maximize visibility of small differences in the generally high correlation values.

**Figure 5.**
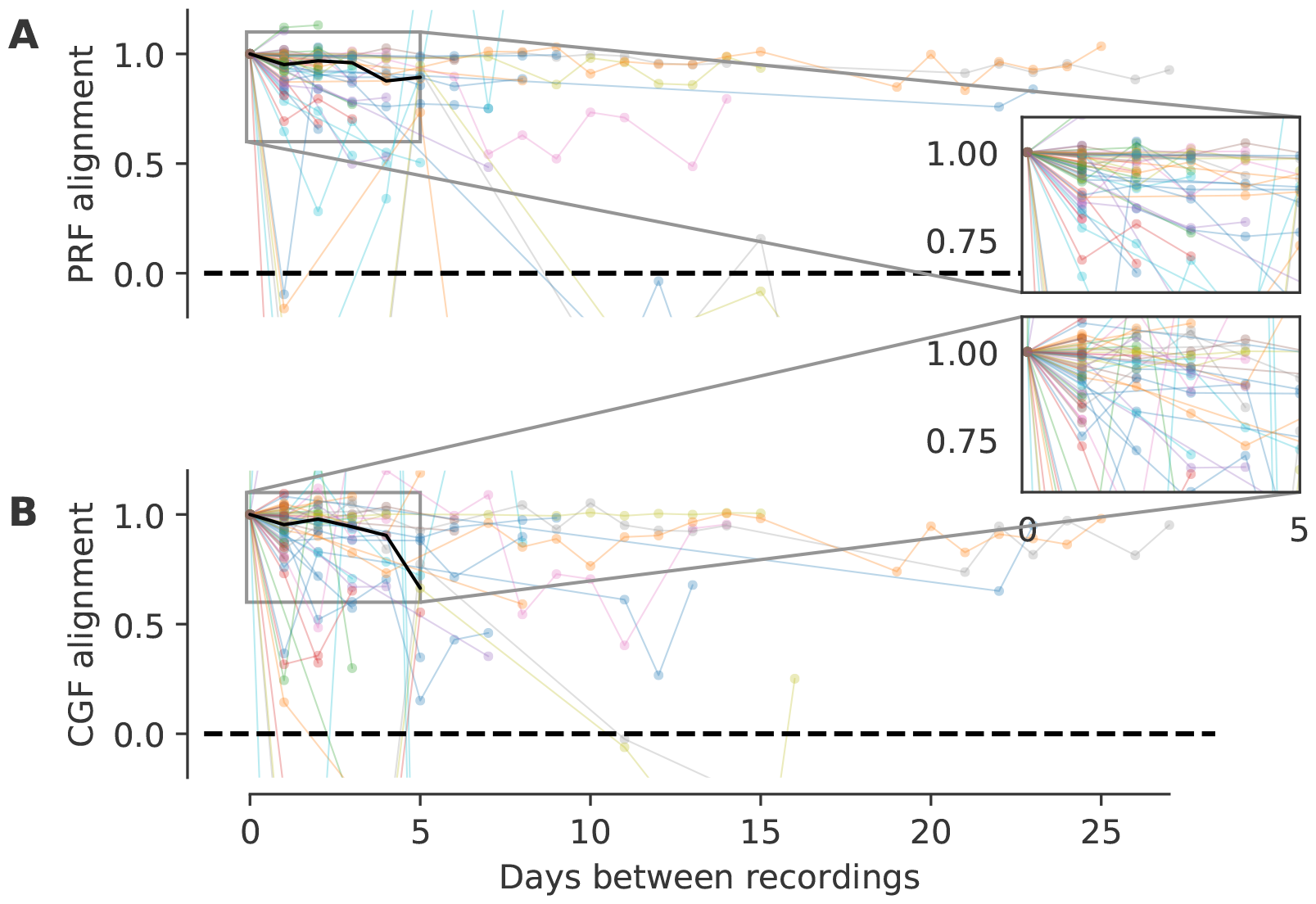
Population data on stability of PRFs and CGFs: normalized field alignment indices. **A-B**. Stability of PRFs (A) and CGFs (B) quantified using a normalized field alignment index, where 1.0 indicates similarity equivalent to the field correlation observed for within-session comparisons for each unit, and 0.0 indicates baseline field correlation expected for comparisons between PRFs or CGFs obtained from different units (see text for details). Data points on day 0 represent the within-session comparison; subsequent points represent comparisons across different numbers of days. Each colored line represents a unit; the solid black line is the median across units. Insets show zoomed-in views of the bulk of the data, between days 0 and 5. Normalized field alignment remained close to 1.0 across sessions for most PRFs and CGFs, indicating that neuron-specific PRF and CGF structure was preserved for many days in most neurons.

To examine stability of PRFs and CGFs across the neuronal population, we first quantified similarity using a normalized *field alignment index*, where 1.0 indicates across-session similarity equivalent to that observed for within-session comparisons for each unit, and 0.0 indicates similarity no higher than the expected baseline for the population (i.e., the similarity that would be expected for comparisons between fields from different units). The field alignment index was calculated as 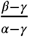,with terms defined as follows for CGFs (and equivalently for PRFs). For each unit, we defined the *within-session similarity α* to be the average correlation between CGFs estimated within the same recording session (i.e., the average across central diagonal values in matrices shown in Figure 4D-E). Likewise, we defined the unit’s *across-session similarity β* for sessions *n* days apart to be the average of the correlation values for all CGF estimates from recordings made *n* days apart (i.e., the average across values in an offset diagonal in matrices shown in Figure 4D-E). Finally, we estimated *baseline similarity γ* by comparing the CGF for the unit to CGFs from other units recorded from the same animal.

Analysis of field alignment indices for the recorded population revealed that most PRFs and CGFs were as stable across days of recording as they were within a single recording session (Figure 5). Indeed, as shown by the extended, nearly horizontal trajectories of some of the colored lines in Figure 5, some units maintained within-session levels of stability in neuron-specific PRF and CGF structure across recording sessions as long as three weeks apart.

Similar stability was evident in analysis of the raw correlation values (i.e., *α* for 0 days between recordings, *β*(*n*) for recordings *n* days apart). For reasons explained in Materials and Methods, we used the raw correlation values rather than the normalized field alignment indices for population analysis of PRF and CGF stability. For both PRFs and CGFs, correlation between fields estimated from the same unit’s responses recorded on different days were typically 0.8-1.0, even when recordings were separated by weeks (Figure 6A-B). To illustrate the dominant trends, Figure 6C and D show the lines of best fit to the field correlation values for each unit’s PRF or CGF respectively, computed as a function of days between recordings using weighted regression. Note that a slope of -0.2 for this best-fit line would correspond to loss of field correlation across 5 days, which was the most common time range over which our repeated recordings were made. As demonstrated in Figure 6E and F, the best-fit line slopes were significantly higher than -0.2 for almost all units (68/69 PRFs, 64/69 CGFs with slope estimates at least 2 standard errors greater than -0.2). Moreover, for both PRFs and CGFs, the slopes were often statistically indistinguishable from zero (29/69 PRFs, 49/69 CGFs with slope estimates within 2 standard errors of 0). Thus, most PRFs and CGFs were stable on a timescale that substantially exceeded the range of our measurements.

**Figure 6.**
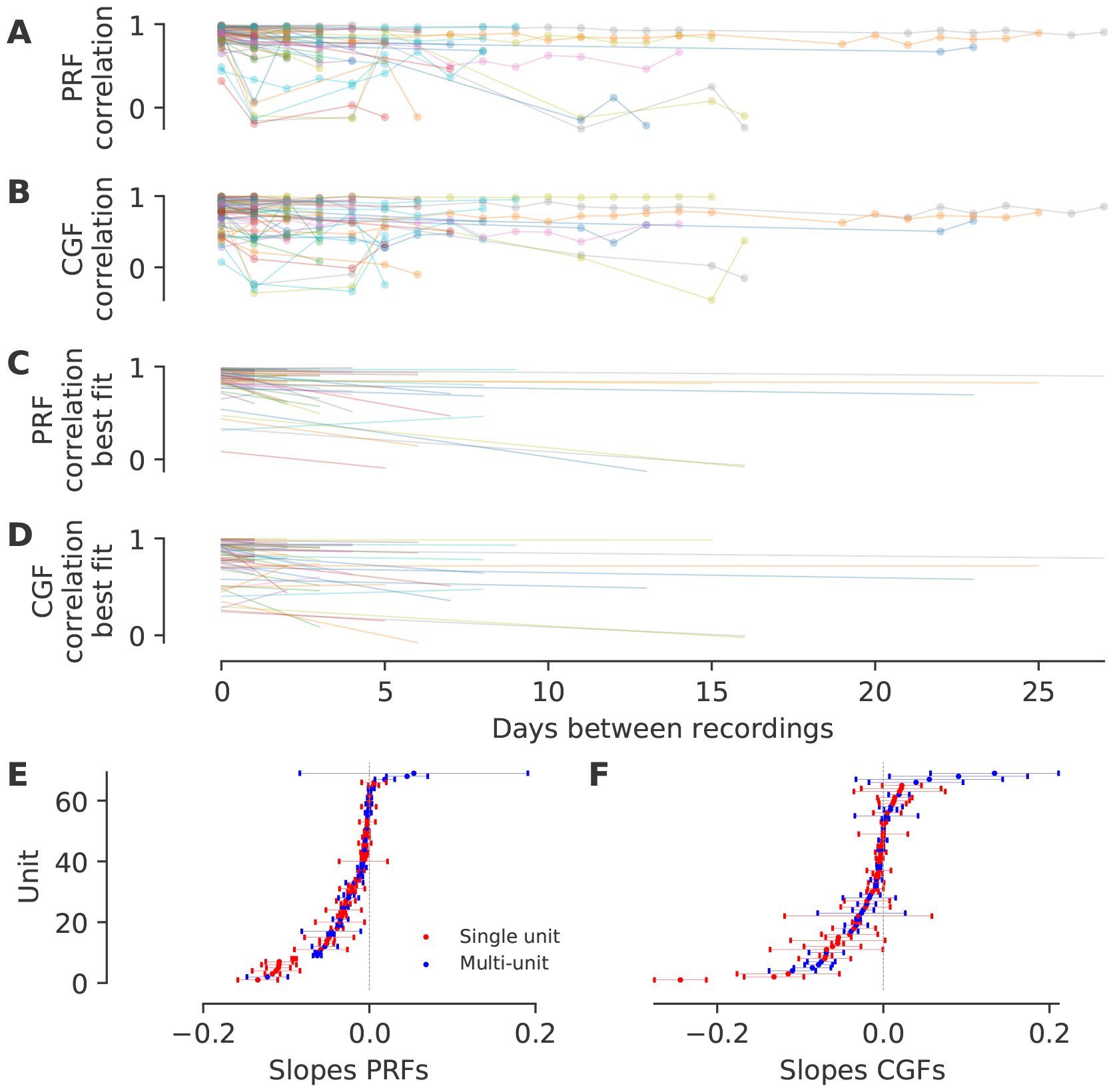
Population data on stability of PRFs and CGFs: raw correlation values. **A-B**. Stability of PRFs (A) and CGFs (B) quantified using raw correlation values, where 1.0 indicates perfect alignment of fields estimated from recordings made on two different days (see Figure 4 for examples). As in Figure 5, data points on day 0 represent within-session comparisons; subsequent points represent comparisons across different numbers of days. Each colored line represents a unit, and lines are transparent so that shading darkens as multiple lines superimpose. Note that most units display high PRF (A) or CGF (B) correlation values that are stable across days or weeks. **C-D**. Lines of best fit to the within-session and across-session field correlation values for each unit, for PRFs (A) and CGFs (B). Each best-fit line was estimated using weighted regression, taking into account the number of within-session (0 days between recordings) and across-session (*n* days between recordings) comparisons available for the unit. **E-F**. Slope (x-axis) for each colored line in A or B respectively; units (y-axis) are ordered by increasing slope. Error bars indicate ±1 standard error of the estimated slope. For both PRFs (C) and CGFs (D), the slopes of the best-fit lines were often statistically indistinguishable from zero and rarely more negative than -0.2 (the value corresponding to loss of field correlation across 5 days).

Further analysis confirmed the conclusion that PRFs and CGFs were both remarkably stable properties of neuronal responses. For example, using the slope of the line of best fit to the field correlation values across recording intervals (Figure 6) as a measure of stability, we found that the distribution of unit-by-unit differences in PRF and CGF stability was strongly peaked at zero (Figure 7A). We also analyzed relationships between the slope measure of PRF and CGF stability and properties of the neuronal response to the DRC stimulus. There was no significant correlation between PRF or CGF stability and either the mean evoked firing rate or the signal-to-noise power ratio of the neuronal response to the DRC stimuli. There was a weak positive correlation between PRF stability and the normalized predictive power of the context model (Spearman’s *rho* = 0.3, *p* = 0.014), but no significant relationship for CGF stability. Thus, PRF and CGF structure appeared to be similarly stable within units and relatively robust to across-unit variation in neuronal response properties or model fits.

**Figure 7.**
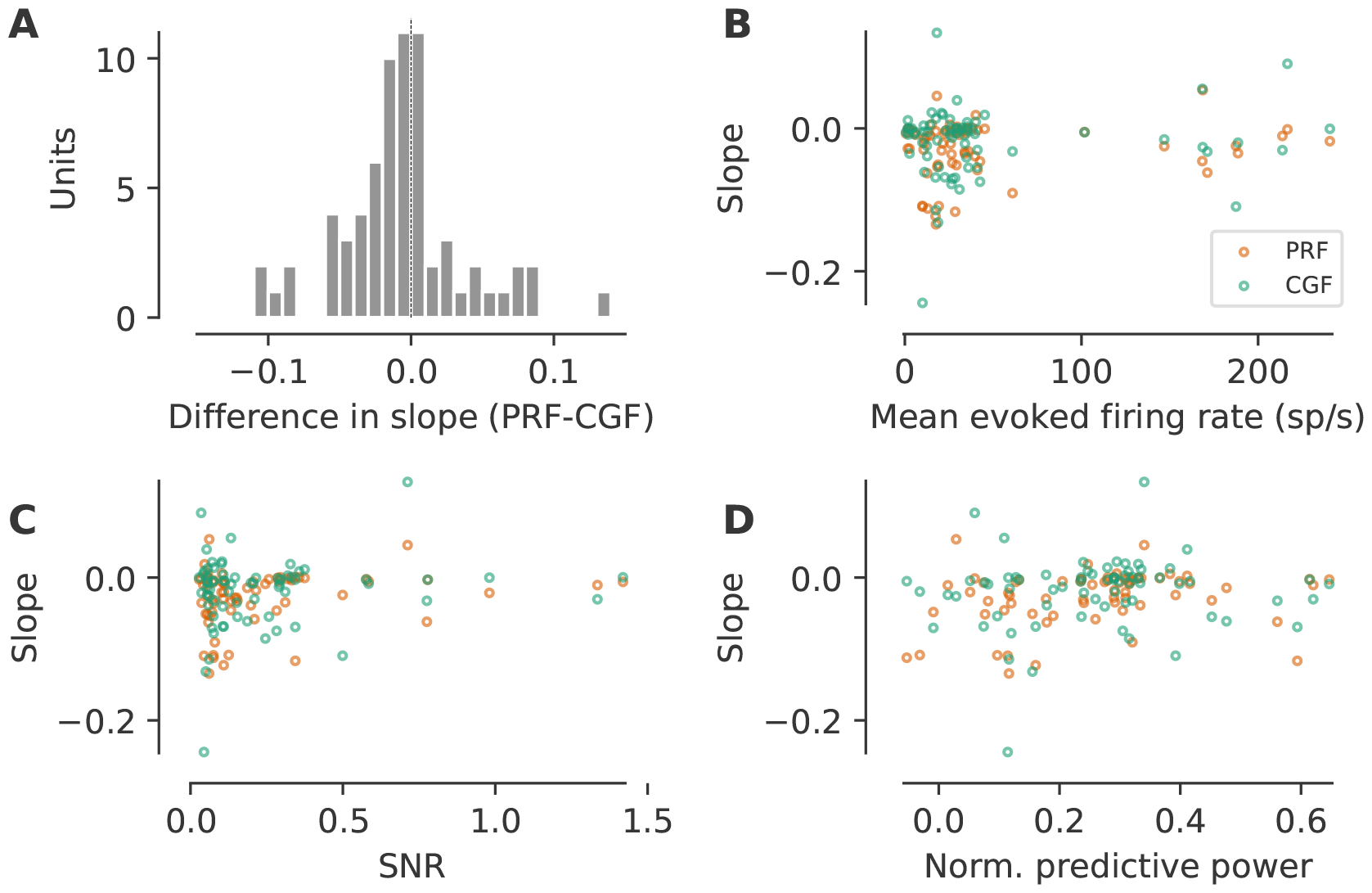
Further analysis of PRF and CGF stability. **A**. Histogram of unit-by-unit differences between the PRF and CGF slope estimates from the correlation-based stability analysis shown in Figure 6. Note clustering of values near zero (dotted line). **B-D**. Slopes of best-fit lines from correlation-based stability analysis for PRFs (orange) and CGFs (green), plotted versus: mean firing rate evoked by the DRC stimulus (B); signal-to-noise power ratio in the neuronal response (C); and normalized predictive power of the context model fit (D). There was no apparent relationship between PRF/CGF stability and firing rate or signal-to-noise power ratio. The stability of PRFs showed a weak positive correlation with normalized predictive power (Spearman’s *rho* = 0.3, *p* = 0.014), whereas that of CGFs did not.

### Changes in locomotion or pupil size have significant but small effects on context model fits to neuronal responses

Given that neuron-specific CGF and PRF structure tended to be stable for days in awake mice, we wondered if there was any effect of the animal’s behavioral state on the context model fits, and by extension, on the spectrotemporal tuning and contextual sensitivities of the neurons. Previous studies have found that excitability of auditory cortical neurons decreases during locomotion (***Schneider et al., 2014***) and either increases with pupil dilation or exhibits non-monotonic dependence on pupil size (***McGinley et al., 2015; Schwartz et al., 2020***). These changes in neuronal excitability with behavioral state are known to modulate the gain and variability of auditory cortical responses to sound, but do not necessarily affect stimulus selectivity (***Schwartz et al., 2020***). We therefore asked whether context model fits might be robust to changes in locomotion and pupil size in awake mice.

We recorded locomotor activity and pupil size along with auditory cortical activity in the vast majority of our experiments (Figure 1), and observed behavioral associations and changes in neuronal excitability consistent with previous reports. For example, in line with previous work (e.g.: ***Reimer et al., 2014; Schneider et al., 2014***), we found that locomotor activity was associated with pupil dilation in mice, and evoked firing rates tended to be smaller when the mouse was moving than when it was still (data not shown).

To investigate how changes in locomotor activity and pupil size affected context model fits, we compared the residuals from model predictions between different behavioral states. Neuronal responses to multiple repetitions of a 45-s-long DRC trial were required to fit each context model, and behavioral variables like locomotion and pupil size varied on a much faster timescale than the DRC trial length (Figure 8A). Hence, it was not feasible to fit different context models to entire DRC trials when the mouse was still versus moving or when the pupil was small versus large. Instead, we analyzed how the moment-by-moment error in context model predictions depended on the animal’s behavioral state. For each unit, we fit a context model to all DRC responses recorded from the cell; calculated the difference between the observed neuronal response and context model prediction for each 20-ms time bin in all DRC recordings; and then compared the interquartile range (IQR) and median of these residuals for time bins when the animal was still versus moving or when the pupil was small versus large (see Materials and Methods for further details).

**Figure 8.**
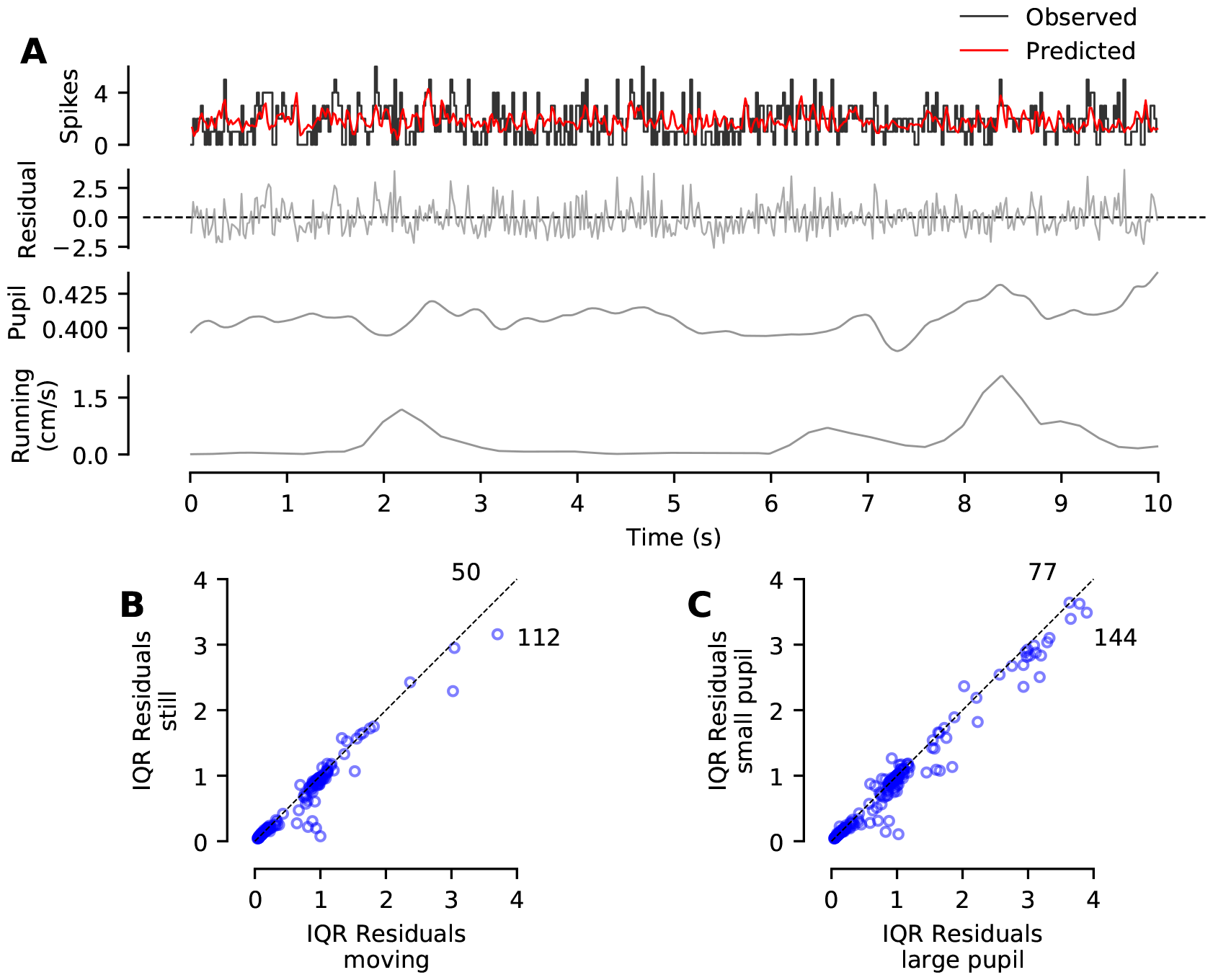
Small but significant effects of locomotor activity and pupil dilation on context model fits. **A**. Observed spiking activity of a single unit (top, black) overlaid with the context model prediction (top, red). Underneath is a trace showing the difference between the two (i.e., the residual, shown in grey, measured in spikes). Dotted black line indicates zero residual. Further below is the pupil diameter (measured as a proportion relative to eye width), and below it a trace of the animal’s running activity over the same period of time. **B**. Interquartile range (IQR) of the residuals for individual units recorded when the mice were still versus moving (i.e., timebins when running cylinder rotation speed was zero versus non-zero). Dotted black line indicates diagonal where the IQRs are equal. Note that more data points fall below than above the line, but that most data points are very close to the diagonal. **C**. Residual IQRs for units recorded when the pupil size was small versus large (i.e., pupil diameter less than or greater than the median pupil diameter for the relevant recording sessions). Conventions and observations as in B.

Analysis of the residual IQRs showed that the ability of the context model to predict temporal variation in auditory cortical responses was significantly, but minimally, affected by changes in locomotor activity and pupil size (Figure 8, B and C respectively). An increase in residual IQR implies poorer prediction of fluctuations in firing rate driven by the DRC stimulus. Comparisons between behavioral states revealed that residual IQRs were significantly higher when the animal was moving or the pupil was large (Wilcoxon sign-rank tests, *p* = 1.5*x*10^−26^ for the locomotion data, *p* = 2.7*x*10^−13^ for the pupil data). However, effect sizes were extremely small in both cases (Cohen’s *d*, 0.09 for the locomotion data and 0.07 for the pupil data). Moreover, the balance of the residual IQRs between the behavioral states improved when the dataset was restricted to single-unit recordings (e.g., proportion of units above the diagonal rose from 0.31 to 0.34 for the locomotion data and from 0.35 to 0.40 for the pupil data). We therefore conclude that behavioral state had significant but very small effects on the ability of the context model to predict temporal fluctuations in auditory cortical responses to the DRC stimulus. In Supplementary Figure 2, we also report small but significant effects of behavioral state on the median residuals from context model predictions; however these results are more difficult to relate to PRF/CGF structure because unlike residual IQRs, median residuals depend not only on the PRF/CGF structure but also on a constant offset term in the model related to prediction of overall mean firing rate.

In sum, these analyses indicate that context model fits were significantly influenced by changes in locomotor activity or pupil size in awake mice, but that effect sizes were small. More detailed analyses of the influence of behavioral state on CGF/PRF structure will require further experiments in which animals are trained or motivated to maintain particular behavioral states for prolonged periods, so that separate context models can be estimated for each state.

## Discussion

Our results indicate that auditory cortical neurons in awake mice maintain remarkably stable patterns of nonlinear sensitivity to combinations of sound input. Individual neurons recorded across many days displayed consistent nonlinear contextual sensitivity (CGF structure) which was stable across many days and comparable in stability to spectrotemporal tuning (PRF structure). In fact, for most auditory cortical neurons, the projected timescale for stability of neuron-specific CGF (and PRF) structure was well beyond the timeframe of the repeated measurements performed here. Average CGF structure in awake mice was qualitatively similar to that observed previously in anesthetized mice (***Williamson et al., 2016***). Notably, however, the ability of the CGF/PRF model to fit temporal fluctuations in the neuronal response was significantly modulated by behavioral state in awake mice, although effect sizes were very small. These observations support the conclusion that both spectrotemporal tuning and nonlinear sensitivity to acoustic context are stable features of auditory cortical receptive fields, which can be at least partially modified by behavioral state. Interestingly, recent two-photon imaging studies in the visual cortex have drawn similar conclusions regarding the stability of orientation tuning, size tuning and surround suppression (***Marks and Goard, 2021; Ranson, 2017***), at least in the highly responsive neurons that would be preferentially sampled with electrophysiological recording techniques. Thus, nonlinear sound-combination sensitivity may well be as stable a feature of auditory cortical receptive fields as orientation selectivity, size tuning and surround suppression are in visual cortical receptive fields.

These results may at first appear in conflict with reports of “representational drift” in the auditory cortex; however, they are not. Recent two-photon calcium imaging studies of auditory cortex in awake mice have concluded that “representational drift” in the auditory cortex arises primarily from fluctuations in the responsiveness of individual neurons, not from changes in the stimulus selectivities of those neurons when they are responsive (***Chambers et al., 2022; Aschauer et al., 2022***). Two-photon calcium imaging allows individual neurons to be tracked over time more definitively than is possible with spike-waveform matching from electrophysiological recording, and it is therefore a better technology to use to address questions about whether individual neurons “drop in” and “drop out” of population activity over time. However, two-photon calcium imaging provides an indirect measure of spiking activity with temporal resolution too low for detailed mapping of receptive fields; therefore, it is limited in its ability to address questions about stability of stimulus selectivity over time. Conversely, while extracellular electrophysiological recording can only be used to track neurons when they are responsive, it allows measurement of spiking activity with the sub-millisecond temporal resolution required for mapping auditory cortical receptive fields. Thus, results of the present electrophysiological study complement and extend the conclusions of previous two-photon imaging studies in auditory cortex, by showing that spectrotemporal receptive fields and sound combination sensitivities of responsive auditory cortical neurons are remarkably stable over time.

To the best of our knowledge, these data also provide the first demonstration that nonlinear sensitivity to acoustic context is a stable, robust feature of receptive fields in the awake auditory cortex. Previous electrophysiological studies of stability in auditory cortical responses to complex sounds have examined consistency of linear spectrotemporal tuning, not nonlinear combination sensitivity, over a timescale of minutes to hours, not days. These studies have reported that spectrotemporal receptive fields (analogous to PRFs, not CGFs) are stable over minutes to hours of recording in awake passively listening animals (***Elhilali et al., 2007; Grana et al., 2009***) and are substantially modified by engagement in behavior (e.g., ***Fritz et al., 2005; David et al., 2012***). Other electrophysiological studies have described input nonlinearities analogous to CGF structure (***Ahrens et al., 2008; David et al., 2009; Pienkowski and Eggermont, 2010; Williamson et al., 2016; Harper et al., 2016; Williamson and Polley, 2019***), but none have examined whether these nonlinearities remain stable over days in individual auditory cortical neurons. We found that both CGFs and PRFs were remarkably stable, with neuron-specific structure that was nearly as consistent between recording sessions conducted on different days as between repeated recordings conducted within the same session on the same day.

These findings have three important implications: physiological, computational, and conceptual. First: physiologically, our results imply that nonlinear sensitivity to acoustic context in the auditory cortex is driven by neural mechanisms that are as stable as those underlying spectrotemporal tuning — at least for sound conjunctions within the <300 ms, ±1 octave range captured by the CGF. Nonlinear contextual sensitivity is sometimes assumed to be a more labile, emergent property of auditory cortex than spectrotemporal tuning, which is largely inherited from strongly stimulus-driven subcortical inputs. However, neurons throughout the subcortical auditory system are known to respond nonlinearly to combinations of sound input. Even within the cochlea, adaptation at the hair-cell synapse generates nonlinear forward suppression in auditory nerve fibres over a ∼50ms timescale, and cochlear mechanics generate nonlinear two-tone interactions spanning many octaves (***Zhang et al., 2001; Zilany et al., 2009***). Other examples of nonlinear combination sensitivity have been documented throughout the auditory brainstem and midbrain, for example in the dorsal cochlear nucleus (***Nelken et al., 1997; Yu and Young, 2000***), the medial nucleus of the trapezoid body (***Englitz et al., 2010***), the nucleus of the lateral lemniscus (***Portfors and Wenstrup, 2001***), and the inferior colliculus (***Portfors and Wenstrup, 1999; Portfors and Felix, 2005; Brimijoin and O’Neill, 2010; Wenstrup et al., 2012***). Finally, previous work has already shown that CGF structure in the auditory thalamus is similar to that observed in the auditory cortex, with small differences in the temporal extent of delayed suppression (***Williamson et al., 2016***). Cortical sensitivity to acoustic context could therefore be predominantly inherited from strongly stimulus-driven, sound-combination-sensitive subcortical inputs — a scenario consistent with both the stability of CGF structure over time and the small effect sizes observed for variation in model fits with spontaneous changes in behavioral state.

Second: computationally, our results call into question common assumptions about sound representation in the auditory cortex — which are implicit in all studies that have used linear spectrotemporal receptive field (STRF) models or linear-nonlinear (LN) models to describe cortical responses to complex sounds (e.g., ***Linden et al., 2003; Depireux et al., 2001; Rabinowitz et al., 2011; Atencio et al., 2008***, and many others). Contrary to the assumptions of STRF and LN models, neurons in the auditory cortex do not linearly combine sound information across points in spectrotemporal space, applying any nonlinear transformations only after the linear combination of sound inputs. Rather, sound inputs are integrated *nonlinearly* across spectrotemporal space, as demonstrated by the robustness and stability of CGF structure. These input nonlinearities cannot be captured by either STRF or LN models. Of course, the nonlinear-linear (NL) CGF/PRF context model also has limitations; for example, it assumes that the same pattern of combination sensitivity applies to all regions of spectrotemporal space (see ***Williamson et al., 2016***, for evidence supporting this assumption), and it does not currently incorporate known output nonlinearities such as variable spiking thresholds. Alternative approaches using nonlinear-linear frameworks have made different assumptions, but reached similar conclusions regarding the relevance of input nonlinearities to complex sound encoding in the auditory cortex (***David et al., 2009; Pienkowski and Eggermont, 2010; Harper et al., 2016; Lopez Espejo et al., 2019***). As famously noted by Box, all models are approximations and therefore wrong, but some are useful (***Box, 1979***). The CGF/PRF model is useful because it reveals that nonlinear combination sensitivity is a robust and stable feature of the neural code for complex sounds in auditory cortical neurons.

Finally, the third and most fundamental implication of our results is that the traditional concept of a sensory “receptive field” is misleading, at least for auditory cortical neurons. A sensory receptive field is typically defined as encompassing the region of sensory space where stimuli evoke changes in the spiking activity of a neuron. Early discoveries in visual neuroscience led to an expansion of this definition to include the concepts of a “classical” (driving) and surrounding “non-classical” (modulatory) receptive field (for a recent review, see ***Angelucci et al., 2017***). However, even this expanded definition of a receptive field is inadequate for describing the robust and pervasive nonlinear sensitivity to sound combinations revealed in the CGFs. Auditory cortical receptive fields are better defined as nonlinear (and therefore context-dependent) filters with stable sensitivities both to individual sensory inputs and to particular combinations of those inputs within a region of sensory space. This alternative conceptualization of a sensory receptive field may be more accurate not only for auditory cortical neurons but also for neurons in other brain areas and sensory systems.

## Materials and Methods

### Animals

A total of 8 male CBA/Ca mice were initially implanted for experiments, and 4 provided sufficient amounts of well-targeted auditory cortex data over months of recording for this study. We used CBA/Ca mice because this strain maintains excellent hearing in adulthood and is therefore a commonly used strain for studies of normal auditory brain function. Following the implantation surgery at 8–12 weeks of age, the mice were singly housed in standard mouse housing rooms, in specially designed cages for mice with implants. Mice were put on a 12-hour reversed light-dark cycle and were provided with food and water ad libitum. Recordings were conducted in each mouse for 3–5 months. All surgical, recording, and housing procedures were performed under a licence approved by the UK Home Office in accordance with the United Kingdom Animals (Scientific Procedures) Act of 1986.

### Chronic tetrode implants

Chronic tetrode implants were custom-made using a microdrive (Axona; UK). A connector (OMNET-ICS; USA) with 34 pins was attached using dental cement. Eight tetrodes each made of 4 strands of 17μm thick platinum 10% iridium wire (California Fine Wire Company; USA) were attached to the connector, with the two remaining pins used for grounding. Tetrodes were plated with platinum to an impedance of 150 kΩ before implantation, and advanced together into the brain using the microdrive.

### Surgery

Mice were chronically implanted with both the tetrode microdrive and a head fixation ring at 8 to 12 weeks of age. The animals were anesthetized with 1.0–3.0% isofluorane and received perioperative and post-operative analgesia (carprofen 5 mg/kg) and post-operative hydration with 0.1 ml warmed saline. A bone screw was inserted in an anterolateral position relative to bregma for a grounding wire. Then, a small craniotomy for the tetrode implant was made over the left hemisphere 2.9 mm lateral and 2.6 mm posterior to bregma. The tetrode bundle was inserted into the brain at an angle of 24^º^to vertical to allow for a roughly tangential microdrive trajectory through the auditory cortex. The initial depth of tetrode bundle insertion along this trajectory was no more than 3.5 mm relative to to the skull surface at bregma; tetrodes were subsequently advanced fully into core auditory cortex using the microdrive after the animal recovered from the surgery. The microdrive, bone screw, and a head fixation ring were secured to the skull using Superbond (C&B Sun Medical; Japan) and dental cement. The grounding wire attached to the bone screw was then soldered to the ground pin on the implant. Following surgery, mice were allowed 2 weeks to recover and acclimatize to head fixation before the commencement of experiments.

### Calibration

As shown in Figure 1, the speaker was directed at the animal’s right ear during experiments, and auditory cortical recordings were made from the left hemisphere. Approximately every month, acoustic stimuli were calibrated with a G.R.A.S. 1/4” microphone positioned where the opening of the animal’s right ear would be during experiments. Calibrations were performed with a G.R.A.S. microphone amplifier and preamplifier (Models 12AA and 26AC). Typically, the calibration ensured that the frequency response of the sound system was flat to within ±2 dB over at least a 5–40 kHz range (more typically, 2–80 kHz). The microphone response was periodically calibrated using a Svantek sound level calibrator emitting a 1 kHz tone at 94 or 114 dB SPL.

### Stimuli

A typical neuronal recording session involved presentation of two identically repeated series of stimuli (called here a “segment”). Each segment consisted of 3 different stimuli separated from each other by 5 seconds of silence. The stimuli were:

- Noise bursts of varying duration and inter-onset timing (used primarily for a separate study and not discussed further here).
- Tone sequences: 10 trials per frequency/intensity combination; tone length 100 ms with 5-ms cosine ramp rise/fall; 20 frequencies equally spaced between 5 kHz and 40 kHz; intensities 40, 50, 60 or 70 dB SPL each with an equal chance; tones of different frequency/intensity combinations presented in a random order.
- Dynamic Random Chord (DRC) stimulus: 15 continuous repetitions of a 45-s-long DRC trial; chords composed of 20-ms tone pips with 5-ms cosine ramp rise/fall; tone pip frequencies 5–40 kHz in 1/12-octave increments; tone pip intensities 25–70 dB SPL in 5 dB increments; average density 6 tones per chord, or 2 tone pips/octave.

See Figure 1 for illustrations of the stimuli and the recording set-up.

### Experimental set-up

Recordings were performed in a sound-attenuating box. Stimuli were generated in Python and converted to analog signals with a sound card (HDSPe AIO; RME; Germany), amplified using a power amplifier (RB-971; ROTEL; Japan) and passed through an attenuator (PA5; Tucker-Davis Technologies; USA) for presentation via a loudspeaker (XT25TG30-04; Peerless Vifa; USA) located approximately 12 cm to the right of the animal’s right ear.

Neural activity was recorded using a 32-channel Intan RHD 2132 amplifier board (hardware bandpass filtering between 1.1 and 7603.8 Hz; Intan Technologies; California, USA), connected to an Open Ephys acquisition board (available from www.open-ephys.org) via an ultra-thin, serial peripheral interface (SPI) cable (RHD2000; Intan Technologies; California, USA). Tetrode recordings were sampled at 30 kHz. The Open Ephys GUI was used to visualize the local field potential and the raw signal was recorded after passing through a bandpass filter of 6-6000 Hz.

A camera was used to record a close-up around the right eye of the mouse, and a second camera recorded the a profile view of the mouse from the right side. Cameras had a sampling frequency of 30 frames/s. Infrared light was added as well as dim visible light in order to keep the pupil diameter at an appropriate level for tracing light-independent changes in pupil size.

The mouse was allowed to run on a custom styrofoam cylinder (20 cm in diameter; 11.5 cm in width; ball-bearing mounted axis) while head-fixed throughout the experiment. A rotary encoder (1024 steps per rotation; Kubler; Germany) was fitted onto the axis of the polystyrene wheel to allow for measurement of the running speed of the mouse during the experiment. Rotation steps were extracted using a microcontroller (Arduino Uno; Farnell; UK) and the running signal was synchronized to neuronal recordings by connecting the microcontroller output onto the Open Ephys data acquisition board.

### Experimental procedures

Mice were accustomed to handling for 2–3 days, acclimatized to the set-up for a further 2–3 days, and finally habituated to increasingly longer periods of head-fixation (from 5 to 30–60 minutes with a daily addition of 5 minutes) in the recording booth. To assess stability of spectrotemporal tuning and contextual sensitivity in auditory cortical neurons across many days, we designed a long experiment where we recorded the neural signal from the same site in the auditory cortex on at least five different (not necessarily consecutive) days, and then advanced the tetrode by 62.5 microns to a new recording site in order to repeat the process (Figure 1C). Locomotor activity and pupil size were continuously recorded along with auditory cortical activity. We recorded from up to 10 recording sites in each mouse. From initial tetrode implantation to the end of recordings, experiments were 3-5 months long for each mouse.

### Data pre-processing

Signals collected from the auditory cortical recordings were spike-sorted using software written in MATLAB (***Sahani, 1999***). The spike-sorting procedure was conducted as follows. First, during the automatic phase, the software would use thresholding to identify potential spikes, then whitened Principal Component Analysis to reduce dimensionality of the data and identify the 4 dimensions which accounted for most of the variance in the waveform shapes. A mixture-of-Gaussians model was then fit to the spike data in the space defined by the 4 principal components. Then, in a later manual phase, the user evaluated the automatically identified clusters to confirm classification as a single unit, multi-unit, or noise, taking into account the following metrics provided by the software.

- The false positive rate and the false negative rate (i.e., the predictions from the mixture-of-Gaussians model for misclassification of spike waveforms inside and outside each cluster). We required that both these rates were below 0.05; otherwise the cluster would be labelled noise.
- Graphical representations of the clusters. Each cluster was represented in 10 different plots (because of the 10 possible combinations between the 4 principal components calculated). If the clusters were not well separated from the noise cluster they were labelled noise. If two or more clusters were superimposed onto each other in all plots, then they were merged.
- The waveform shapes for the spikes of each cluster. In rare cases the user would re-classify as noise an automatically identified “unit” cluster for which the waveform shape seemed biologically implausible. For example, if the waveform shape had qualities reminiscent of electrical noise or if it seemed inconsistent between spikes, or if it looked exactly the same across the four channels, then the user would take this into account in deciding how to label the cluster.
- Auto-correlograms and cross-correlograms comparing the firing of any two clusters. Cross-correlograms were helpful in identifying clusters that required merging with one another. Auto-correlograms allowed us to decided whether clusters representing genuine neural activity were in fact single units or multi-unit activity. Clusters were labelled as single units when the autocorrelogram showed no firing in the first 2-ms bin after the cluster had just fired. Otherwise, the cluster would be labelled as multi-unit activity.

### Identifying core auditory areas

Following conventions used previously in other studies of mouse auditory cortex (including ***Williamson et al., 2016***), we imposed two physiological criteria in order identify recording sites in “core” auditory areas (i.e., primary auditory cortex and anterior auditory field):

1. A significantly altered firing rate in the first 50 ms following a tone presentation, based on a Wilcoxon test conducted between the firing rates of the unit in interest in the 50 ms before and after stimulus onset in each trial presented. Only cases with significant differences (*p* < 0.01) were deemed core auditory cells.
2. A response latency smaller than 20 ms. The latency was marked as the first 2-ms bin where the response of the cell deviated from the mean spontaneous firing rate (as estimated from the 50 ms before stimulus presentation) by 3 or more times the standard deviation of the spontaneous firing rate.

Units recorded from the same tetrode and same location at the same time as as another unit which directly met these criteria were also designated as core auditory. Therefore, consistent with the approach used in most studies of auditory cortex, the core auditory dataset included both units that met the classic physiological criteria themselves, and neighbouring units that did not.

### Spike-waveform matching

We used a spike-waveform-matching technique previously described in ***Tolias et al. (2007***) to match units found in discontinuous recordings from the same sites. We compared each unit to all units identified on the same tetrode and at the same depth.

Let us consider two example tetrode waveforms to be compared to each other: *X* and *Y* . *X* was first scaled by a factor *α* so as to minimize the sum of squared differences between the two waveforms. Two different metrics were then computed (here referred to as *d*_1_ and *d*_2_). *d*_1_ is the normalized Euclidean distance between the scaled waveforms which therefore represents a metric of the difference in shape that exists between the two waveforms:

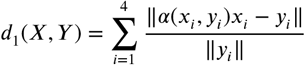

where *α*(*x, y*) = *arg*_*α*_*min* ‖ *αx* − *y*‖^2^, and where the sum is over the four channels of the tetrode. *d*_2_ is defined as:

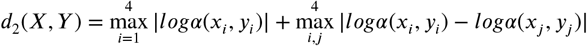

and it captures the difference in the amplitudes across the four channels. As in ***Tolias et al. (2007***), in order to make these measures symmetric we used *d*_1_(*X, Y*) + *d*_1_(*Y, X*) for the shape factor and *d*_2_(*X, Y*) + *d*_2_(*Y, X*) for the scaling factor.

We established a null distribution which consisted of comparisons between units identified on the same tetrode but which were recorded from locations at least 250 μm apart from each other (i.e.: units which were definitely not matches). To compare the two distributions in one dimension we fitted a mixture of two Gaussians model (m) to the experimental data:

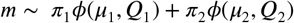

where μ_*i*_ and *Q*_*i*_ represent the mean and covariance matrix of the *i*^th^ component which is normally distributed.

The parameters of one of the two Gaussians in the model were fixed to those which described the null distribution obtained earlier, and the others were learnt using an Expectation-Maximization algorithm. This second Gaussian therefore describes the distribution of possible matches. We then took as confirmed matches the waveform comparisons which fell within the 99% confidence interval of the distribution of possible matches and outside the 99% confidence interval of the null distribution. Using these stringent criteria we aimed to minimize the false positives, and maximise the chances that matched recordings truly originated from the same unit.

### Context model fitting

For context model analysis, we required that core auditory units met three additional physiological criteria related specifically to their responses to the DRC stimulus.

- Each unit’s DRC responses had to have signal power at least one standard error greater than zero. Context models can be fit effectively only to units which show stimulus-dependent variation in their responses to the DRC stimulus (i.e., non-zero signal power).
- Firing rate over all trials had to be greater than or equal to 5 spikes/s. When neurons fire very infrequently this leads to highly sparse matrices which create issues with the fitting of the model.
- Normalized noise power had to be below below 40. This was a criterion put in place to remove outliers with excessive noise power.

Mathematically, the context model generates predictions for the response (*r*) of a neuron to a given sound (*s*) with spectrotemporal energy at time t in frequency channel f, as follows:

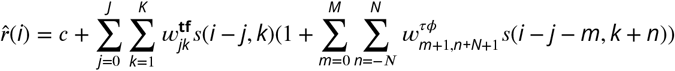

where the constant *c* sets a baseline firing rate, *w*^**tf**^ is the PRF field with summation limits over time shifts (t) and frequency (f), and *w*^*τϕ*^ is the CGF field and the summation limits are over relative time shift (*τ*) and relative frequency (*ϕ*). The CGF weight corresponding to zero time-frequency offset was fixed to 0; therefore no weight contributed to its own context, resulting in a linear STRF model prediction for presentations of isolated tones. For more details on the model and validation of its performance, see ***Williamson et al. (2016***).

The context model was fitted for each unit to the neuronal responses recorded during DRC segments. Typically we had 2 DRC stimuli of 15 trials each, per recording session, for each unit. The first DRC trial from each segment was discarded to minimize effects of level adaptation on context model fits. The fitting was done using Alternate Least Squares (ALS) as there were two fields whose parameters needed to be optimized (the PRF and the CGF). We used Automatic Smoothness Determination (ASD) (***Sahani and Linden, 2002a***) to find the best smoothing parameters for the PRFs. We then fixed those smoothing parameters for the PRF and then used a grid search method to find the best smoothing parameters for the CGF, because the ASD method did not reliably converge when both PRF and CGF smoothing were simultaneously optimized.

The grid search method involved running multiple fittings of the model with the PRF smoothing parameters fixed and each time changing the combination of the three smoothing parameters for the CGF. At the end we proceeded with the CGF smoothing parameters that provided the greatest cross-validation predictive power. The parameters we tried during the grid search method were the following: scaling parameter: (3, 6 and 9); spectral smoothing parameter: (2, 4, 6, 8, 10, 12) and temporal smoothing parameter: (1.5, 3.5, 5.5, 7.5, 9.5). These values were chosen for the grid search to try and cover as much parameter space as possible, but were centered around the parameter values that were most commonly observed in CGFs.

To ensure that differences in smoothing parameters did not confound assessments of field stability for each unit, we estimated optimal smoothing parameters for the context model for each unit using all the data available for the unit (i.e., all DRC recordings made across days). These optimal smoothing parameters for the unit were then applied to all context model fits based on individual DRC segments.

### Stability assessment

To evaluate similarity between fields (PRFs or CGFs) estimated in different DRC segments, we calculated the normalized dot product (i.e., zero-shift two-dimensional cross-correlation) between the fields, without subtraction of the mean matrix. The within-session field correlation was simply the correlation between the two fields estimated from the first and second DRC segment of a session respectively. The across-session field correlation was the average correlation across all four possible comparisons between the two DRC segments in each of the two recording sessions. We then constructed a similarity matrix for each unit which shows on the central diagonal the within-day, within-session field correlations and on the offset diagonals the across-day field correlations. The method is illustrated graphically in Figure 1D and for example neurons in Figure 4D-F.

For each unit, we then defined the within-session field similarity *α* to be the average withinsession field correlation for CGF (or PRF) estimates from all the unit’s recording sessions. Likewise, the across-session similarity *β* for sessions *n* days apart was defined as the average of all the unit’s across-session field correlation values for CGF (or PRF) estimates obtained from recordings made *n* days apart. We plotted these within-session and across-session field similarity values versus the number of days between recording sessions, and estimated the best-fit line to the data using weighted linear regression (taking into account the number of comparisons contributing to the averages *α* and *β*(*n*)). We used the slope of this best-fit line as our measure of the unit’s CGF (or PRF) stability for population analysis. This slope represents the average rate of change in CGF (or PRF) correlation as a function of time between recording sessions, but does not necessarily imply a gradual rate of change across chronological days of recording. It should be noted that similar slopes could be obtained from a gradual decline in field correlation over time and from a more abrupt drop on a particular day during the recording period for a unit.

We also examined CGF (and PRF) stability using a normalized field alignment index, where 1.0 represents within-session similarity for the unit and 0.0 indicates baseline similarity expected for comparisons with CGF (or PRF) estimates from other units. For each unit’s CGF (or PRF), we defined the field alignment index as 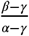, where *α* and *β* were calculated as above, and *γ* was a baseline field correlation measure obtained by comparing the CGF (or PRF) for the unit to those from other units recorded from the same animal. This normalized field alignment index was useful for comparing across-session to within-session stability and assessing persistence of neuron-specific CGF (or PRF) structure (Figure 5). However, unlike the raw field correlation values and field similarity measures *α* and *β*, the values of the normalized field alignment index were potentially unbounded and very noisy in units with poor within-session stability. For this reason, we used the field similarity measures rather than the normalized field alignment indices to derive the slope estimates used for population analyses of CGF and PRF stability (Figure 6 and Figure 7).

### Pupil diameter extraction

For pupil tracking we used DeepLabCut (version 2) (***Mathis et al., 2018; Nath et al., 2019***), which leverages a ResNet-50-based convolutional neural network for predicting the location of a desired bodypart across frames. We labeled 80 frames taken from 4 videos (one from each animal), then used 95% of the video data for training. We trained for 1,030,000 iterations, validated with 1 shuffle and we report a test error of 3.38 pixels and a train error of 1.03 pixels (image size was 416 by 252). We then used a p-cutoff of 0.99 to condition the predicted coordinates, which served to exclude predictions for which the network was not certain, hence making our data more reliable. We used this network to analyze videos collected under very similar experimental conditions. During labelling, we marked 8 specific points: 6 points delineating the edges of the pupil and 2 at the left and right edges of the mouse’s eye.

We fit an ellipse on the pupil in each frame. We asserted that there had to be at least 5 out of 6 points in a frame in order for us to attempt to fit the ellipse. The major axis of the ellipse (in pixels) fitted to the pupil in that frame was then taken as the diameter of the pupil. Some videos were excluded after manual inspection due to poor quality of tracking.

We estimated the width of the eye opening during a recording as the median distance between the points marking the right and the left edge of the eye, which the network was also trained to detect. Finally, in our analyses we used the frame-wise pupil diameter, obtained as explained above and normalized by the median length of the eye in each recording, meaning that our measurements are not influenced by small perturbations in the positioning and angling of the camera relative to the animal’s eye from day to day of experimentation.

### Analysis of effects of locomotor activity and pupil dilation

To assess the effects of locomotor activity and pupil dilation on context model performance we divided the timebin-by-timebin data for each unit into categories of still versus moving, or small versus large pupil. More specifically, for the locomotor activity analysis, we divided all the 20-ms timebins from each recording with locomotor activity data into two categories: (i) timebins when the mouse was still (speed = 0.0; 36.0% for the average unit, standard deviation 15.1%), or (ii) timebins when the speed of the mouse was non-zero (62.7% of timebins for the average unit, standard deviation 16.3%). Results of the residuals analysis were similar when we used a higher speed threshold for the still/moving categorization (e.g., with threshold 0.5 cm/s, 83.3% timebins categorized as still, 15.3% as moving, standard deviation 12.5% and 11.6% respectively). For the pupil size analysis, we divided all the timebins into two categories: (i) timebins when the pupil was more dilated than the median pupil size for all sessions for the unit, or (ii) timebins when the pupil was smaller than the median. Since this categorization was performed relative to the median pupil size, equal percentages of timebins fell into the two categories.

We then calculated the residual from the context model prediction of the firing rate in each timebin (observed minus predicted firing rate), and computed the interquartile range and median of the residual distributions for the different timebin categories for each unit. The IQRs and medians for the residual distributions with and without locomotor activity or pupil dilation were then compared across units in population analysis (Figure 8 and Supplementary Figure 2). Note that we used the actual signed residual rather than the absolute residual, in order to distinguish cases in which the observed firing rates were either higher or lower than predicted. Further interpretation of these measures is provided in the main text.

## Acknowledgments

This work was supported by the Biotechnology and Biological Sciences Research Council (BB/P007201/1, JFL); the London Interdisciplinary Doctoral Programme (MA); the Simons Foundation (SCGB543039, MS); and the Gatsby Charitable Foundation (MS). We would also like to thank Simon Rumpel for constructive discussions which helped us to improve this manuscript. These discussions were enabled by a workshop supported by grant NSF PHY-1748958 to the Kavli Institute for Theoretical Physics (KITP).

## Supplementary Figures

**Supplementary Figure 1.**
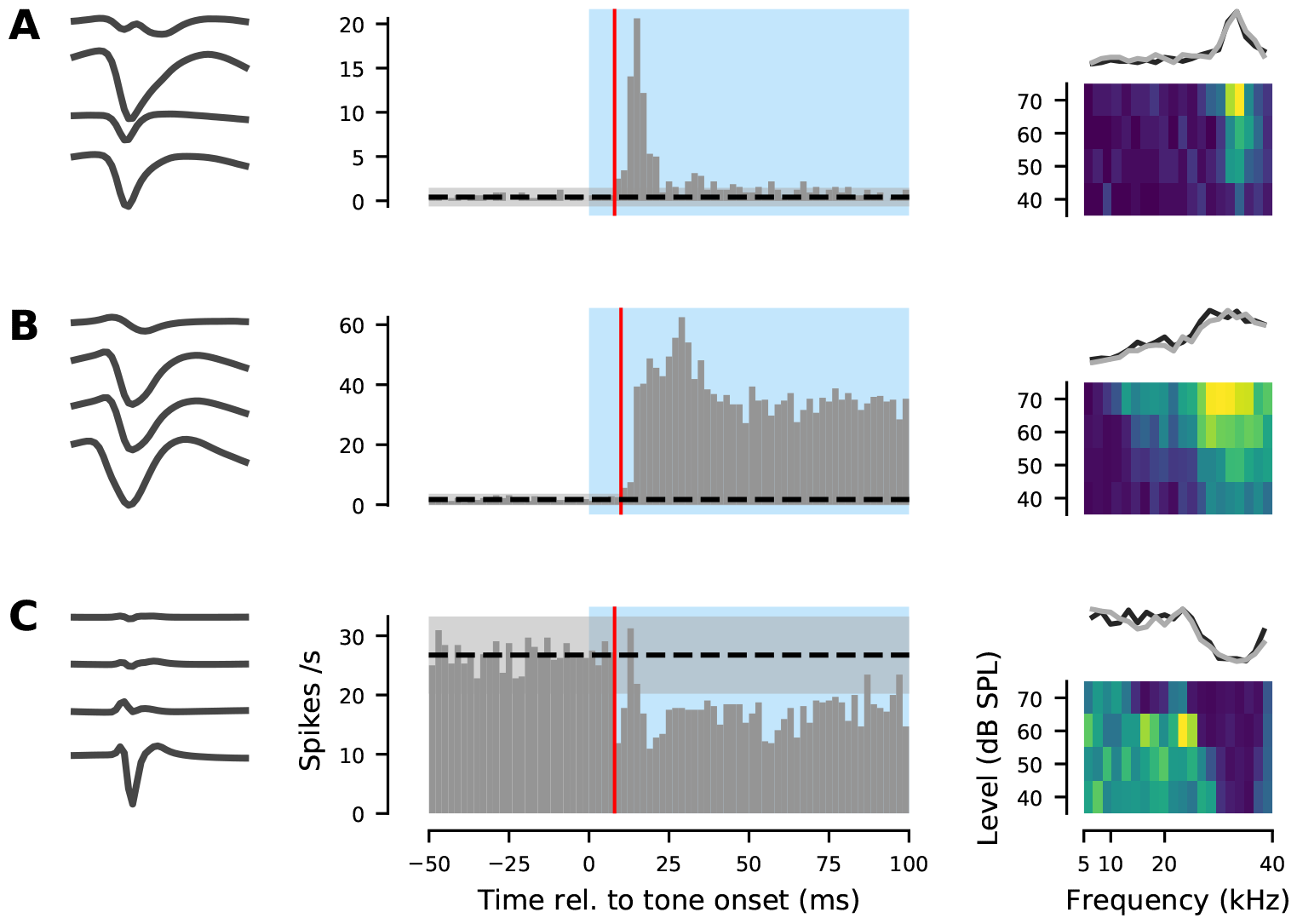
Core auditory cortical recording sites identified using physiological criteria. **A-C**. Three example units recorded from one animal. Left: raw waveform on each of the four tetrode channels. Middle: PSTH of the response of each unit to 100-ms pure tones, pooled across tone frequencies and intensities. Blue shading indicates time of tone presentation. All three units met the two criteria for classification as “core” auditory cortex: (1) robust responses to tone pips (significant difference in firing rate across trials between the 50 ms before and 50 ms after tone onset; Wilcoxon rank-sum test, *p* < 0.01), and (2) response latency <20 ms. Latency is indicated here with a red vertical line and was defined as the first time bin after tone onset where the firing rate fell outside the mean (dotted black line) ±3 standard deviations (grey shaded area) of the bin-by-bin firing rates in the 50 ms before tone onset. Right: Frequency-Response Areas (FRA). Top of each panel: frequency tuning profile averaged over all tone intensities. The grey and black lines indicate estimates of the frequency tuning profile obtained from two different runs of the tone-pip sequence separated by more than 20 minutes. The overlap of these two lines illustrates the consistency of frequency tuning estimates in units with the robust, short-latency responses typical of “core” auditory cortex.

**Supplementary Figure 2.**
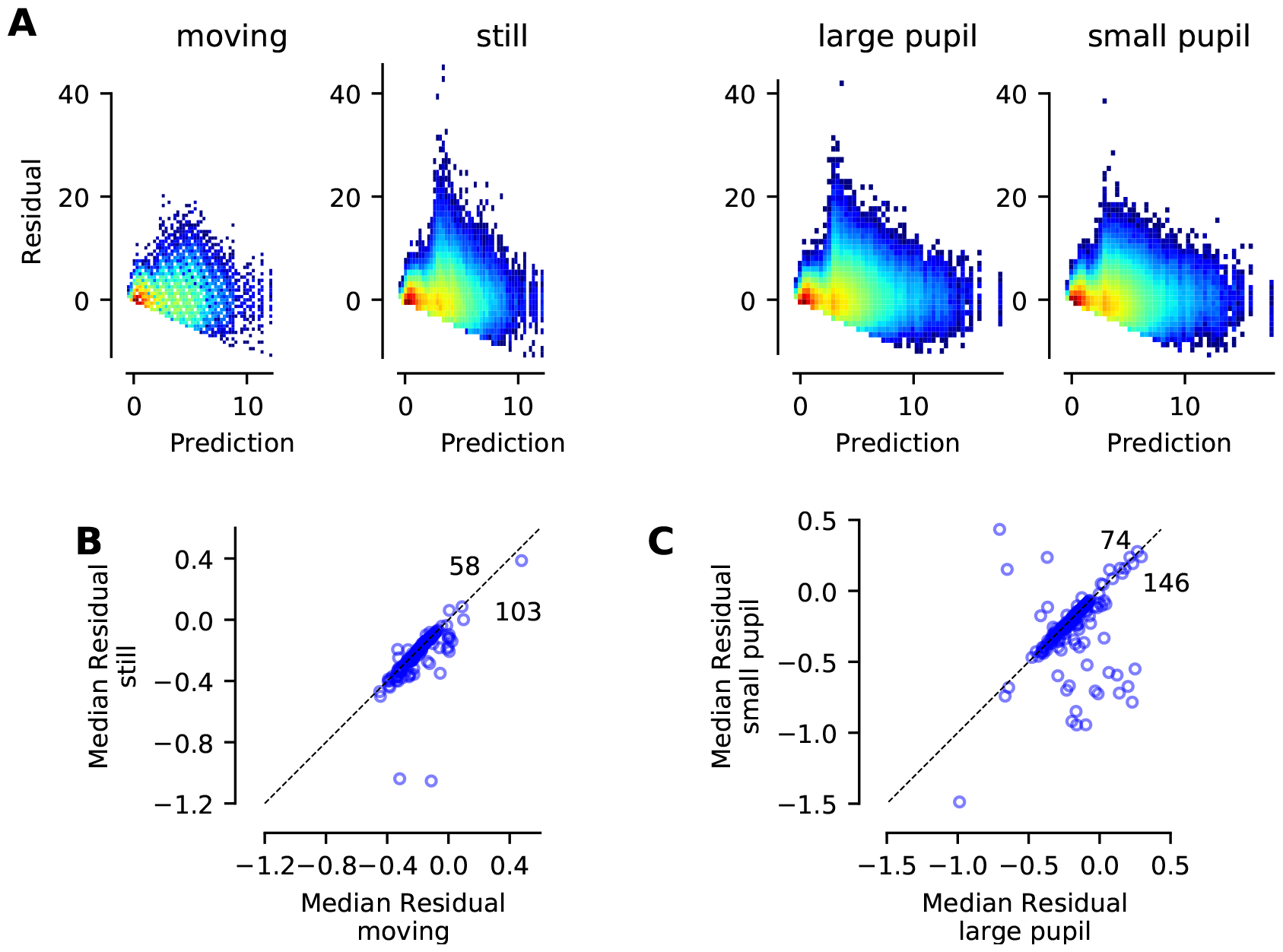
Effects of locomotor activity and pupil dilation on context model residual distributions and median residuals. **A**. 2D histograms of the context model residuals plotted against the model predictions when mice had: (i) non-zero speeds (leftmost plot); (ii) zero speed (still: middle left); (iii) a pupil diameter above the median diameter for the unit recordings (middle right); and (iv) a pupil diameter below the median diameter (rightmost plot). Pupil median diameters were calculated for each unit based on all data available from the relevant recording sessions. Plots show pooled data from all suitable timepoints in recordings from all mice. **B-C**. Scatter plots showing the median of the residuals when the mice were still versus moving (B) or when the pupil size was small versus large (C). Dotted black line indicates equal values. Note that median residuals were typically slightly negative, indicating that the context model tended to over-predict firing rates. Note also that median residuals were significantly more positive (i.e., in most cases, less negative) when the animal was moving or the pupil was large (Wilcoxon sign-rank tests: locomotion data, *p* = 1.8*x*10^−24^; pupil data, *p* = 3.7*x*10^−14^). Effect sizes were relatively small (Cohen’s *d*: locomotion data 0.23; pupil data 0.29), but not as tiny as for residual IQRs.

